# Modulatory effect of *Syzygium aromaticum* and *Pelargonium graveolens* on oxidative stress and inflammation

**DOI:** 10.1101/362426

**Authors:** Ilias Marmouzi, El Mostafa Karym, Rachid Alami, Meryem El Jemli, Mourad Kharbach, Fouzia Mamouch, Bouchra Faridi, Aisha Attar, Yahia Cherrah, My El Abbes Faouzi

## Abstract

**Background:** Therapy combination is defined as disease treatment with two or more medication to acheive efficacy with lower doses or lower toxicity. Regarding its reported toxicities and efficacy, the Essential Oils (EOs) from Syzygium aromaticum (SA) and *Pelargonium graveolens* (PG) were combined for *in vitro* and *in vivo* assays and toxicities.

**Methods:** The Essential Oils and mixture were tested for *in vivo/in vitro* antioxidant and anti-inflammatory activities. The assays included the animal model of acute inflammation (carrageenan model), the protective effect on H_2_O_2_/Sodium nitroprissude induced stress in *Tetrahymena pyriformis*, and the *in vitro* antioxidant assays.

**Results:** The chemical analysis of the investigated Oils has lead to the identification of Eugenol (74.06%), Caryophyllene (11.52%) and Carvacrol acetate (7.82%) as the major element in SA; while PG was much higher in Citronellol (30.77%), 10-epi-γ-Eudesmol (22.59%), and Geraniol (13.95%). In our pharmacological screening of samples, both Oils demonstrated good antioxidant effects. *In vivo* investigation of the antioxidant activity in the protozoa model (*T. pyriformis*) demonstrated a lesser toxic effect of EOs mixture with no significant differences when oxidative stress markers and antioxidant enzymes (MDA, SOD and CAT) were evaluated. On the other hand the *in vivo* model of inflammatory response to carrageenan demonstrated a good inhibitory potential of both EOs. The EOs Mixture demonstrated equivalent bioactivity with lower toxic effect and minimal risk for each compound.

**Conclusions:** The results from this study indicate that EOs mixture from SA and PG demonstrated promising modulatory antioxidant/anti-inflammatory effect, which suggest an efficient association for therapy.

## 1. Introduction

Oxidative stress and inflammation are closely related processes, simultaneously found in many pathophysiological conditions such as diabetes, hypertension, cardiovascular and neurodegenerative diseases [1-2]. Therefore, antioxidant therapy could be of great interest in treating inflammatory diseases. However, the failure of antioxidant trials might result from the selection of inappropriate agents for both inflammation and oxidative stress or the use of nonselective agents that block some of the oxidative and/or inflammatory pathways but exaggerate the others [3]. In this regards, classical phytotherapy and herbal combinations is known for higher efficacy and lesser side effects. Therefore any progress or optimization of plant synergy may lend to standardized formulations with multitarget focus. This has the advantage of reducing side-effects due to the lower doses of active compounds within the extract mixtures.

Essential Oils (EOs) are a mixture of several compounds characterized by an essence of aromatic plants. Approximately 3000 EOs are known, from which 300 are commercially used in the pharmaceutical, cosmetic and food industries, as well as complementary therapies [4]. Many of those EOs components (such as Eugenol, Citronellol and Geraniol) are known for their anti-inflammatory and protective effect against oxidative stress. In fact, the phenolic compound Eugenol have shown many disparencies in data literature, for instance Fonsêca et al., [5], reported that Eugenol reduces leukocyte migration and inhibits COX-2 without affecting COX-1. While Paula Porto [6], revealed that Eugenol do not change COX-2, NF-κB1 or TNF-α expression. On the other hand, Eugenol might modulate inflammatory processes and DNA damage or induce primary DNA lesions depending on the concentration. According to Teho Koh [7], Eugenol show bi-modal actions, by stimulating or inhibiting the IL-8 production at lower and higher concentrations. Then, it is clearly noticed that Eugenol at high concentrations appears to exert cytotoxic effects, while low concentrations are able to produce anti-inflammatory activity via NF-κB inhibition; In fact those variations are related to Eugenol phenoxyl radicals [8-9]. Another interesting compound is the monoterpene alcohol Citronellol that inhibits carrageenan neutrophil migration and iNOS enzymatic activity and attenuate COX-2 and LPS-induced mRNA expression. Citronellol similarly to Eugenol shows a dose–response relationship phenomenon characterized by low-dose stimulation and high-dose inhibition [10]. Finally Geraniol an acyclic monoterpene alcohol was found to significantly decrease lipid peroxidation, inhibit NO release and ROS generation. Abe et al., [11], demonstrated that Geraniol clearly suppressed TNF-α induced neutrophil adherence. Moreover, Geraniol inhibits murine skin tumorigenesis by modulating COX-2 expression, Ras-ERK1/2 signaling pathway and apoptosis (Sandeep Chand Chaudhary). Veerasamy Vinothkumar et al., [12], found that Geraniol modulates cell proliferation, apoptosis, inflammation, and angiogenesis during 7,12-dimethylbenz[a]anthracene-induced hamster buccal pouch carcinogenesis.

Despite the interesting anti-inflammatory and antioxidant activities of those EOs compounds, their bi-modal action and easy switch from beneficial effect to toxic manifestations let us suggest the choice of a multi-extracts combination for fewer side effects with an equivalent or enhanced pharmacological properties. In this regards two EOs known for their interesting levels of these compounds were chosen for our experiments. *S. aromaticum* which contains a high percentage of Eugenol [13], and *P. graveolens* which contains considerable amounts of Citronellol and Geraniol [14-15].

## 2. Materials and Methods

### 2.1. Ethical considerations

*P. graveolens* was harvested at the botanical garden of faculty of medicine and pharmacy in Rabat in May 2015 (No specific permits were required for the described field studies or for the collection of plant material). *S. aromaticum* cloves has been purchased from the market in Rabat, and used for further investigations.

Ethics approval for animal experimentations was obtained from Mohammed V University in Rabat. The experiments were conducted in accordance with the accepted principles outlined in the “Guide for the Care and Use of Laboratory Animals” prepared by the National Academy of Sciences and published by the National Institutes of Health and all efforts were made to minimize animal suffering and the number of animals used.

### 2.2. Phytochemical analysis

The crude protein content was determined using the Kjeldahl method. Crude fat was determined by extracting a known aliquot of sample (100 g) with petroleum ether, using a Soxhlet apparatus. Acid detergent fibre (ADF), lignin content (ADL) and cellulose were also determined. The mineral composition (Ca, Fe, K, P, Na, Zn, Se, Mg and Mn) was determined using an inductively coupled plasma atomic emission spectroscopy (ICP AES, Jobin Yvon Ultima 2). The amount of phenolic contents (PC) was determined according to the Folin-Ciocalteu method. While, the flavonoids contents (FC) in the fractions were determined using a colorimetric method. The phenolic content was determined as mg of Gallic acid (mg GAE/g EOw) equivalent per g of Essential Oil weight. All procedures were described previously [16].

### 2.3. Essential Oils extraction

The Essential Oil was extracted from *P. graveolens* and *S. aromaticum* by hydro-distillation using Clevenger apparatus. The distillations were carried out on 500 g for 4 hours, after which the water content of Oil was eliminated using anhydrous K_2_CO_3_.

### 2.4. Gas chromatography-mass spectrometry

Gas chromatography combined with mass spectrometry (GC-MS) was used for the identification of the main compounds. The analysis was performed on a GC-MS Clarus 600/560DMS PerkinElmer (Bridgeport, USA), equipped with an automatic injector. The system is controlled by mass Turbo Software (Windows XP SP2). The stationary phase is a column supelco^®^ (L 30m x 0.25 ID x DF 0.25) Elite-5MS phase, the carrier gas is helium at a flow rate of 0.8mL/min. The oven was programmed from 75 °C to 320 °C at a gradient of 10 °C per minute with a pre-heating the transfer line at 325 °C and 250 °C at source. The automatic injector in splitless is guide (50/1 to 250 °C). Ionization is caused by electron impact (EI). Identification of the extracts components was based on computer matching with NIST and Wiley 275 libraries.

### 2.5. In vitro antioxidant properties

The free radical scavenging activity of the essential oils was measured by 2,2’-Diphenyl-1-picrylhydrazyl hydrate (DPPH). The radical-scavenging activity was calculated as a percentage of DPPH discoloration and represented as trolox equivalent from the standard curve. The ABTS+ cation radical was produced by the reaction between 10 mL of 2 mM ABTS in H2O and 100 μL of 70 mM potassium persulphate, stored in the dark at room temperature for 24 h. The antioxidant activities samples are expressed as TEAC values, defined as the concentration of standard trolox with the same antioxidant capacity of the fraction under investigation (mg TE/g EOw). Moreover, the ferric ions (Fe^3+^) reducing antioxidant power (FRAP) method was used to measure the reducing capacity of Essential Oils. The reducing power of the Oils was represented as ascorbic acid equivalent (mg AAE/g EOw). Antioxidant assays procedures were described previously [16].

### 2.6. Protective effect against oxidative and nitroprusside stress

#### Microorganism and cell culture

The *T. Pyriformis* strain was maintained axenically in 5 mL of proteose-peptone (1.5 %, w/v) and yeast extract (0.25 %, w/v) (PPYE) medium at room temperature (25 ± 1 °C). Cultures of *T. Pyriformis* were prepared in 500 mL erlenmeyer flasks containing 100 mL of sterile PPYE medium inoculated with 1% (v/v) of 72 hour pre-culture, in the same medium, grown at 32 °C without shaking.

#### Oxidative and nitroprusside stress

The ciliated protozoan *T. Pyriformis* was cultivated over 140 hours in PPYE medium containing H_2_O_2_ (Fluka) or sodium nitroprusside (SNP) (Sigma) at 50 % inhibitory concentrations (IC_50_). These concentrations were determined from *T. Pyriformis* growth inhibitory curves of the two stress reagents. The IC_50_ values were then estimated using prohibits analysis (respectively 0.7 and 1.8 mM for H_2_O_2_ and SNP). Cell growth was monitored microscopically by counting cell numbers at different time intervals using a haemocytometer (Malassez cell). A control was performed in the same conditions without the stress reagents.

#### Essential Oils cytotoxicity

The assay to determine minimum inhibitory concentration (MIC) of the EOs studied was performed by the two-fold serial dilution method. The EO was dissolved in dimethylsulfoxide (DMSO) and then a dilution series from 10^−1^ to 10^−6^ were prepared. Five microliters of each dilution was added to 5 mL of PPYE medium in the test tube. Then, 50 μL of a calibrated inoculum (1.5 - 105 cells/mL) of T. Pyriformis was added to each tube that contained Oil. A tube containing only DMSO (0.1%, v/v) was used as control. The MIC results were taken as the lowest concentration of the EO that showed no turbidity after 72 hours of incubation at 32°C. Each EO was assayed in triplicate.

#### Stress markers under treatment

To evaluate the antioxidant potential of the EOs, *T. Pyriformis* was cultivated in the presence of the stress reagent at the IC_50_ (0.7 mM for H2O2 or 1.8 mM for SNP) in PPYE medium supplemented with the EO tested at a non-toxic concentration of 10^−9^. The stress reagent and the EO were added simultaneously to the growth medium before inoculation. Growth curves were performed and control experiments were carried out using only the PPYE medium inoculated with *T. Pyriformis* in presence of the stress reagent. The protein content was estimated following the method of Lowry. The antioxidant enzymes were expressed as μmol/min/mg protein. Catalase (CAT) activity was determined according to the method of Aebi. T-SOD activity was determined by the method of Beauchamp and Fridovich. Thiobarbituric acid reactive substances were estimated as malondialdehyde (MDA) equivalent (nmol MDA/mg protein), from the calibration curve.

### 2.6. Anti-inflammatory activity

The carrageenan-induced paw edema model was used to evaluate the anti-inflammatory effect of EOs. The initial paw volume was recorded using an UgoBasile model LE750 plethysmometer. Rats groups were orally administered by EOs and mixture at 250 mg/kg; Indomethacin (10 and 20 mg/kg) or distilled water (5 mL/kg). Carrageenan (0.05 mL of a 1% w/v solution, prepared in sterile saline) was injected subcutaneously into subplantar region of the left hind paw of each rat 30 minute later. The paw volumes of each rat were measured at three time points (90, 180 and 360 minutes). Mean differences of treated groups were compared with the mean differences of the control group. The percentages of inhibition of inflammation were calculated according to the following formula:

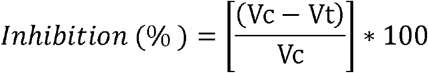

Where Vc is the Mean increase in paw volume of control group; Vt is the increase in paw volume of animals in treated groups. Besides, area under the curve (AUC) was calculated.

### 2.7. Statistical analysis

The data are presented as mean ± SEM (standard error of mean) of six mice. Statistical analysis of the data was performed using one-way analysis of variance (ANOVA) followed by Tukey posthoc test. Significant differences were set at P values lower than 0.05.

## 3. Results

### 3.1. Phytochemical and mineral composition

Table 1 shows selected nutrients compositions of PG and SA. Proteins content (%) in PG (19.12 ± 0.98) was much higher from SA (9.41 ± 0.34). On the other hand, SA content of fat (10.75 ± 1.10 %) and essential oil (5.60 ± 0.67 %) seem to be much interesting than its equal in PG (2.30 ± 0.91 % and 0.12 ± 0.03 %). Phenolic content (Table 1) varied from 8.12 mg GAE/g EO in PG to 27.56 mg GAE/g EO in SA. The high phenolic content in SA is due to the presence of the phenolic acid Eugenol in high concentrations (74.06 %).

**Table 1.**
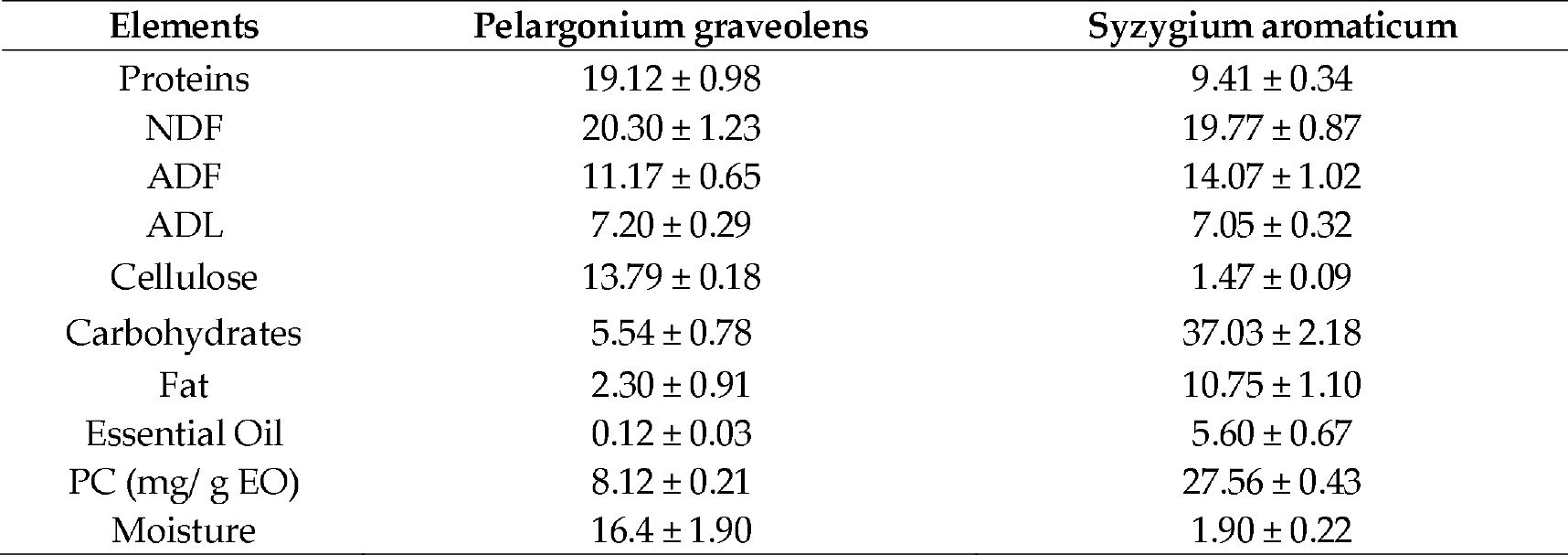
Proximate analysis (%)

| Elements | *Pelargonium graveolens* | *Syzygium aromaticum* |
| --- | --- | --- |
| Proteins | $19.12 \pm 0.98$ | $9.41 \pm 0.34$ |
| NDF | $20.30 \pm 1.23$ | $19.77 \pm 0.87$ |
| ADF | $11.17 \pm 0.65$ | $14.07 \pm 1.02$ |
| ADL | $7.20 \pm 0.29$ | $7.05 \pm 0.32$ |
| Cellulose | $13.79 \pm 0.18$ | $1.47 \pm 0.09$ |
| Carbohydrates | $5.54 \pm 0.78$ | $37.03 \pm 2.18$ |
| Fat | $2.30 \pm 0.91$ | $10.75 \pm 1.10$ |
| Essential Oil | $0.12 \pm 0.03$ | $5.60 \pm 0.67$ |
| PC (mg/ g EO) | $8.12 \pm 0.21$ | $27.56 \pm 0.43$ |
| Moisture | $16.4 \pm 1.90$ | $1.90 \pm 0.22$ |

Mean contents of each mineral found in both plants expressed in mg/kg of dry weight are shown in Table 2. Significant differences (p<0.05) have been registered in the amount of minerals between SA and PG. SA is higher in potassium (K) (10750.90 ± 198.89), magnesium (Mg) (2610.56 ± 32.12), manganese (Mn) (1150.47 ± 77.76), sodium (Na) (7500.39 ± 119.87), and phosphorus (P) (1130.73 ± 36.32). While PG is higher in Selinium (Se) (70.11 ± 9.34) and calcium (Ca) (9450.98 ± 120.23). Iron (Fe) (140.87 ± 2.12) and Zinc (Zn) (210.31 ± 19.90) content was comparable between both plants.

**Table 2.** Mineral contents (mg/kg)

| Minerals | *Pelargonium graveolens* | *Syzygium aromaticum* |
| --- | --- | --- |
| Ca | $9450.98 \pm 120.23$ | $5990 \pm 109.21$ |
| Fe | $140.87 \pm 2.12$ | $130.33 \pm 4.12$ |
| K | $1700.12 \pm 32.34$ | $10750.90 \pm 198.89$ |
| Mg | $740.13 \pm 16.43$ | $2610.56 \pm 32.12$ |
| Mn | $10.07 \pm 0.12$ | $1150.47 \pm 77.76$ |
| Na | $3320.22 \pm 55.32$ | $7500.39 \pm 119.87$ |
| P | $330.34 \pm 11.23$ | $1130.73 \pm 36.32$ |
| Se | $70.11 \pm 9.34$ | $20.98 \pm 2.14$ |
| Zn | $210.31 \pm 19.90$ | $180.67 \pm 8.56$ |

### 3.2. Essential Oils composition

Quantitative analyses of the chemical composition of the investigated EOs have been performed using GC-MS. GC-MS analysis revealed the presence of 18 elements in SA and 5 elements in PG EO. Chemical identification of the Oil constituents was conducted based on their retention time (Rt) and mass spectral data. It was also accomplished using computer search of mass spectral databases. The identified compounds in each EO account for 82.02 % in PG and 99.37 % in SA (Table 3). Pelargonium graveolens EO was mainly composed of Citronellol (30.77%), 10-epi-γ-Eudesmol (22.59 %), Geraniol (13.95 %), Geranyl tiglate (7.48 %) and Geranyl propanoate (7.23 %). Syzygium aromaticum EO was particularly rich in Eugenol (74.06 %), Caryophyllene (11.52%) and Carvacrol acetate (7.82%).

**Table 3.** Essential Oils composition (%)

| N° | Compounds | Formula | PG | SA | Mixture* |
| --- | --- | --- | --- | --- | --- |
| 1 | Furfural | $C_5H_4O_2$ | - | 0.06 | 0.03 |
| 2 | 2-Heptanone | $C_7H_{14}O$ | - | 0.14 | 0.07 |
| 3 | 2-Heptanol, acetate | $C_9H_{18}O_2$ | - | 0.19 | 0.09 |
| 4 | β-Ocimene | $C_{10}H_{16}$ | - | 0.09 | 0.04 |
| 5 | 2-Nonanone | $C_9H_{18}O$ | - | 0.16 | 0.08 |
| 6 | Benzyl acetate | $C_9H_{10}O_2$ | - | 0.17 | 0.08 |
| 7 | Ethyl benzoate | $C_9H_{10}O_2$ | - | 0.25 | 0.12 |
| 8 | Methyl salicylate | $C_8H_8O_3$ | - | 1.19 | 0.59 |
| 9 | Citronellol | $C_{10}H_{20}O$ | 30.77 | 0.40 | 15.58 |
| 10 | Nerol 2,6-Octadien-1-ol, 3,7-dimethyl-, (Z)- | $C_{10}H_{18}O$ | - | 0.36 | 0.18 |
| 11 | Geraniol 2,6-Octadien-1-ol, 3,7-dimethyl-, (E)- | $C_{10}H_{18}O$ | 13.95 | - | 6.97 |
| 12 | Citronellyl formate | $C_{11}H_{20}O_2$ | - | 0.27 | 0.13 |
| 13 | Anethole | $C_{10}H_{12}O$ | - | 0.19 | 0.09 |
| 14 | Eugenol | $C_{10}H_{12}O_2$ | - | 74.06 | 37.03 |
| 15 | β-Elemene | $C_{15}H_{24}$ | - | 0.05 | 0.02 |
| 16 | Carvacrol acetate | $C_{12}H_{16}O_2$ | - | 7.82 | 3.91 |
| 17 | Caryophyllene | $C_{15}H_{24}$ | - | 11.52 | 5.76 |
| 18 | Humulene | $C_{15}H_{24}$ | - | 1.95 | 0.97 |
| 19 | Geranyl propanoate | $C_{13}H_{22}O_2$ | 7.23 | - | 3.61 |
| 20 | Caryophyllene oxide | $C_{15}H_{24}O$ | - | 0.50 | 0.25 |
| 21 | 10-epi-γ-Eudesmol | $C_{15}H_{26}O$ | 22.59 | - | 11.29 |
| 22 | Geranyl tiglate | $C_{15}H_{24}O_2$ | 7.48 | - | 3.74 |
| Monoterpene hydrocarbons |  |  | - | 0.09 | 0.04 |
| Oxygenated monoterpenes (MO) |  |  | 51.95 | 85.26 | 68.56 |
| Sesquiterpenes hydrocarbons (SH) |  |  | - | 13.52 | 6.75 |
| Oxygenated sesquiterpenes (SO) |  |  | 30.07 | 0.5 | 15.28 |
| Total |  |  | 82.02 | 99.37 | 90.63 |

### 3.4. *In vitro* antioxidant activities

There are many assays used for the determination of antioxidant activity of EOS. In the present work, three complementary methods were used to assess the antioxidant activity of SA, PG and EOs mixture: DPPH and ABTS free radical scavenging, and ferric-reducing power assays (Figure 1). SA EO displayed higher antioxidant activity across the three testing methods (DPPH, ABTS and FRAP with 150.19 ± 4.73 mg TE/g EO; 110.86 ± 2.29 mg TE/g EO; 34.82 ± 0.14 mg AAE/g EO) compared to PG (8.12 ± 0.19 mg TE/g EO; 1.85 ±0.22 mg TE/g EO; 19.23 ± 0.07 mg AAE/g EO). The higher antioxidant potential of SA EO might be explained partly by its high content of Eugenol. The antioxidant activity of this phenol has been previously reported [18-19-20], and strong positive correlation was found between antioxidant activity and the content of Eugenol [21]. Oxygenated monoterpenes and sesquiterpenes, particularly Citronellol, Geranial and 10-epi-γ-Eudesmol, the main compounds of P. graveolens are also characterized by their powerful antioxidant capacity, partially attributed to the presence of strongly activated oxygen. The functional groups in those compounds [21-23], might explaine the potent antioxidant activity of P. graveolens essential oil. This result supports the idea that these volatile compounds have a lower antioxidant activity than that of Eugenol [18-24].

**Figure 1.**
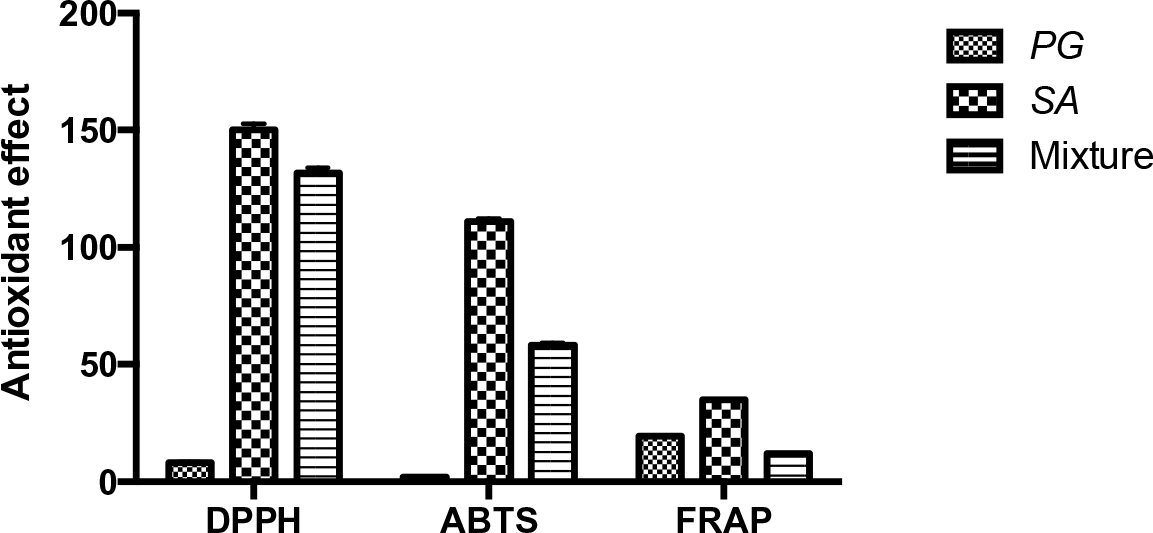
Antioxidant activities of Essential Oils and mixture. DPPH (mg TE/g EO), ABTS (mg TE/g EO), FRAP (mg AAE/g EO)

Comparing the results of antioxidant activity of SA and PG in the present work with data from previous studies, it was found that SA from Brazil and from China possesses an excellent anti-radical effect using the ABTS assay [21]. Also, it appears that SA from Grabowo Wielkie exhibited a strong antioxidant activity using the DPPH, ABTS and FRAP assays [24],. Many authors have reported antioxidant and radical scavenging properties of PG EOs [26-27-28]. However, to our knowledge, the antioxidant effect of EOs combinations has not been previously reported. The EO mixture showed intermediate antioxidant activity and was able to reduce the stable, purple-coloured radical DPPH into yellow-coloured DPPH-H with 131.66 ± 2.46 mg TE/gEO. Similarly, in ABTS and reducing power assays, oils mixture exhibited a considerable antioxidant activity, with 58.16 ± 0.95 TE/g EOand 12.02 ±0.01 TE/gEO, respectively.

### 3.5. Protective effect against stress agents in *T. Pyriformis*

#### 3.5.1. EO cytotoxicity

The EOs can be anti-protozoal agent [29], This activity is caused by the presence of cytotoxic compounds like, terpenes which disrupts the cell membrane [30]. The correct approach before using EOs consist to evaluate the toxicity and to determine the minimum inhibitory concentration (MIC value). It was evident that pure EOs were highly cytotoxic. Therefore, the use of a series of dilution was made to determine the MIC values. The MIC value obtained for PG and mixture was 10^−4^ and MIC value for SA Oil was 10^−6^. For further experiments, we did use the concentration of 10^−9^ which has no toxic effect.

#### 3.5.2. EO protective effect against oxidative stress

For in vitro Oxidative stress induction on the protozoan *T. pyriformis*, we did apply a range of concentrations between 0.3 and 3 mM of H_2_O_2_. After 24 hours of culture, cell growth was assessed and the results (Figure 2) show that H_2_O_2_ has a dose-dependent inhibition of cell growth.

**Figure 2.**
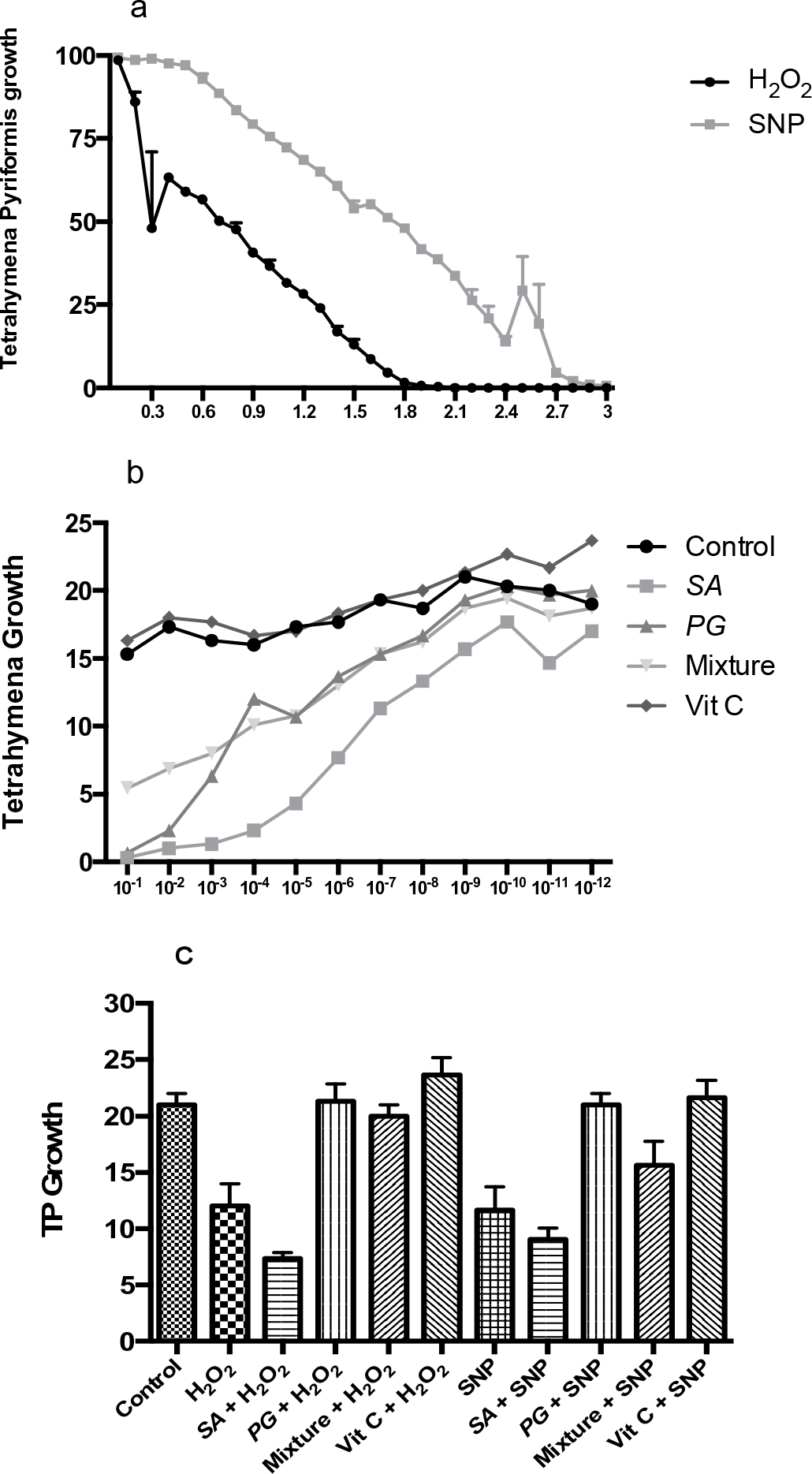
*In vivo* antioxidant activity. a) cytotoxic effect of stress agents. b) Cytotoxic effect of Essential Oils. c) *T. pyriformis* growth under stress and treatements.

The concentration of 1.8 mM completely inhibits cell growth. In these conditions the IC50 of H_2_O_2_ was deduced as 0.7 mM. Evaluation of stress markers showed that H_2_O_2_ (0.7 mM) is a good stress agent that induces lipid peroxidation and increases the enzymatic activity of catalase and SOD. H_2_O_2_ induced stress reduced *T. pyriformis* viability by 40.00% compared to the control group.

Cell viability in treated groups was interestingly improved by 70.00% in PG, 60.00% in the mixture and 78.38% in Vc compared to H_2_O_2_ group; on the other hand SA reduced the cellular growth by −22.50 % compared to H2O2 group. SOD levels increased significantly by 70.29% in H_2_O_2_ compared to the non-treated control (p<0.05). However Oil treatment reduced SOD release by 69.19% in PG, 44.62% in the mixture and 40.16% in the positive control (Vc); and increased its levels by 8.28% in SA. Similarly, the CAT activity in stressed control increased significantly (103.25%) compared to the control. Treatment with EOs and vitamin C decreased CAT levels by 4.06% in SA, 101.62% in PG, 95.11% in mixture and 88.89% in Vc. Treatment of cells with EOs shows that SA Oil shows low protective effect against H_2_O_2_, but the use of PG or the mixture protects cells against the harmful effects of H_2_O_2_.

In our experimental conditions, the evaluation of oxidative stress markers shows that the H_2_O_2_ increases the activities of catalase and SOD. These enhanced antioxidant defenses support a generation of oxygen free radicals in the presence of H_2_O_2_. SOD and catalase, which converts superoxide anions into H_2_O_2_, and H_2_O_2_ into (H_2_O + ½ O2), respectively, probably increase in order to prevent the production of reactive oxygen species, and to reduce their associated side effects: lipid peroxidation, protein carbonylation, and DNA damages contributing to cell dysfunctions leading to cell death [31]. H_2_O_2_ is also a substrat of catalase. [32], the enhancement of catalase activity could also be a consequence of the increased intracellular level of H_2_O_2_. The antioxidative properties of SA, PG and the mixture of EOs observed on *T. pyriformis* could be explained by the chemical properties revealed by antioxidant assays: DPPH, FRAP and ABTS.

#### 3.5.3. Oil protective effect against nitroprusside stress

The nitrosative stress induction is achieved *in vitro* by the effect of SNP on the protozoan *T. pyriformis*. We used a range of concentration of SNP. After 24 hours of culture, cell growth is assessed by direct counting using hemocytometer on optical microscope. Our results (Figure 2a) show that SNP Dose-dependent inhibition of cell growth. The concentration of 3 mM completely inhibits cell growth. Evaluation of stress markers showed that (1.8 mM) is a good stress concentration that induces lipid peroxidation and increases the enzymatic activity of catalase and SOD. SNP induced stress reduced *T. pyriformis* viability by 55.55% compared to the control group Treatment of cells with EOs shows a protective effect, especially of PG and EO mixture (Table 4).

**Table 4.** Stress markers in *Tetrahymena pyriformis* under treatments.

| TP | MDA | SOD | CAT |
| --- | --- | --- | --- |
| **Normal groups** |  |  |  |
| Control | 98.66 ± 0.57 | 1405.00 ± 78.07 | 123.00 ± 2.64 |
| VC | 75.33 ± 1.52 | 1398.00 ± 11.00 | 109.00 ± 2.00 |
| SA | 103.33 ± 2.08 | 2513.00 ± 61.53 | 230.33 ± 2.15 |
| PG | 63.33 ± 4.16 | 1414.66 ± 5.50 | 120.66 ± 1.15 |
| Mixture | 83.33 ± 4.16 | 1648.00 ± 11.26 | 156.66 ± 3.78 |
| **Oxidative stress** |  |  |  |
| H2O2 | 123.00 ± 2.64 | 2392.66 ± 169.77 | 250.00 ± 1.00 |
| Vc + H2O2 | 87.66 ± 2.51 | 1430.33 ± 23.35 | 140.66 ± 3.05 |
| SA + H2O2 | 104.66 ± 2.51 | 2509.33 ± 8.62 | 245.00 ± 1.00 |
| PG + H2O2 | 84.00 ± 4.00 | 1420.33 ± 26.08 | 125.00 ± 3.00 |
| Mixture + H2O2 | 106.66 ± 2.51 | 1765.66 ± 50.08 | 133.00 ± 4.00 |
| **Sodium Nitroprusside stress** |  |  |  |
| SNP | 99.00 ± 1.00 | 1824.33 ± 52.77 | 124.33 ± 3.21 |
| Vc + SNP | 78.66 ± 1.52 | 1396.00 ± 4.58 | 111.66 ± 2.51 |
| SA + SNP | 107.33 ± 2.08 | 2497.33 ± 11.93 | 236.00 ± 3.60 |
| PG + SNP | 81.66 ± 3.21 | 1408.66 ± 18.01 | 125.66 ± 3.21 |
| Mixture + SNP | 92.66 ± 2.08 | 1515.00 ± 8.88 | 157.33 ± 2.51 |

T-SOD and CAT are expressed in $\mu\text{mol}/\text{mg}$ protein. Thiobarbituric acid reactive substances were estimated as malondialdehyde (MDA) equivalent (nmol MDA/mg protein). Vc (Vitamin C).

### 3.6. Anti-inflammatory effect

The anti-inflammatory activity of EOS has been evaluated by carrageenan-induced rat paw edema method. The results are shown in Figure 3. At the evaluated dose (250 mg/kg), the EOs exhibited significant (p < 0.05) inhibitions of increase in paw edema from 1h30 min to 6h after injection of carrageenan as compared to the control group. The edema development of the control group was 48.05 % at 1h30 after carrageenan injection and remained more or less constant till the end of 6h. However, the edema development of the treated group with 250 mg/kg of PG and 250 mg/kg of SA EOs at 1h30 after carrageenan injection were 18.90% and 12.77% respectively and were progressively reduced to 12.68% and 7.49% at 3h and to 13.01% and 3.45 % at 6h respectively. Based on percentages of inhibition values, the combination of both EOs showed a considerable anti-inflammatory activity with 86.17%, 87.79% and 84.84 % inhibition respectively produced at 1h30, 3 h and 6h after the experimental induced paw edema. This effect was comparable to that produced by 10 mg/kg and 20 mg/kg of indomethacin (IND) at 1h30, 3 h and 6h after the experimental induced paw edema with respectively 71.17%, 86.89 %, 78.59 - 83.94 % and 67.63 - 75.63 % of inflammation inhibition.

**Figure 3.**
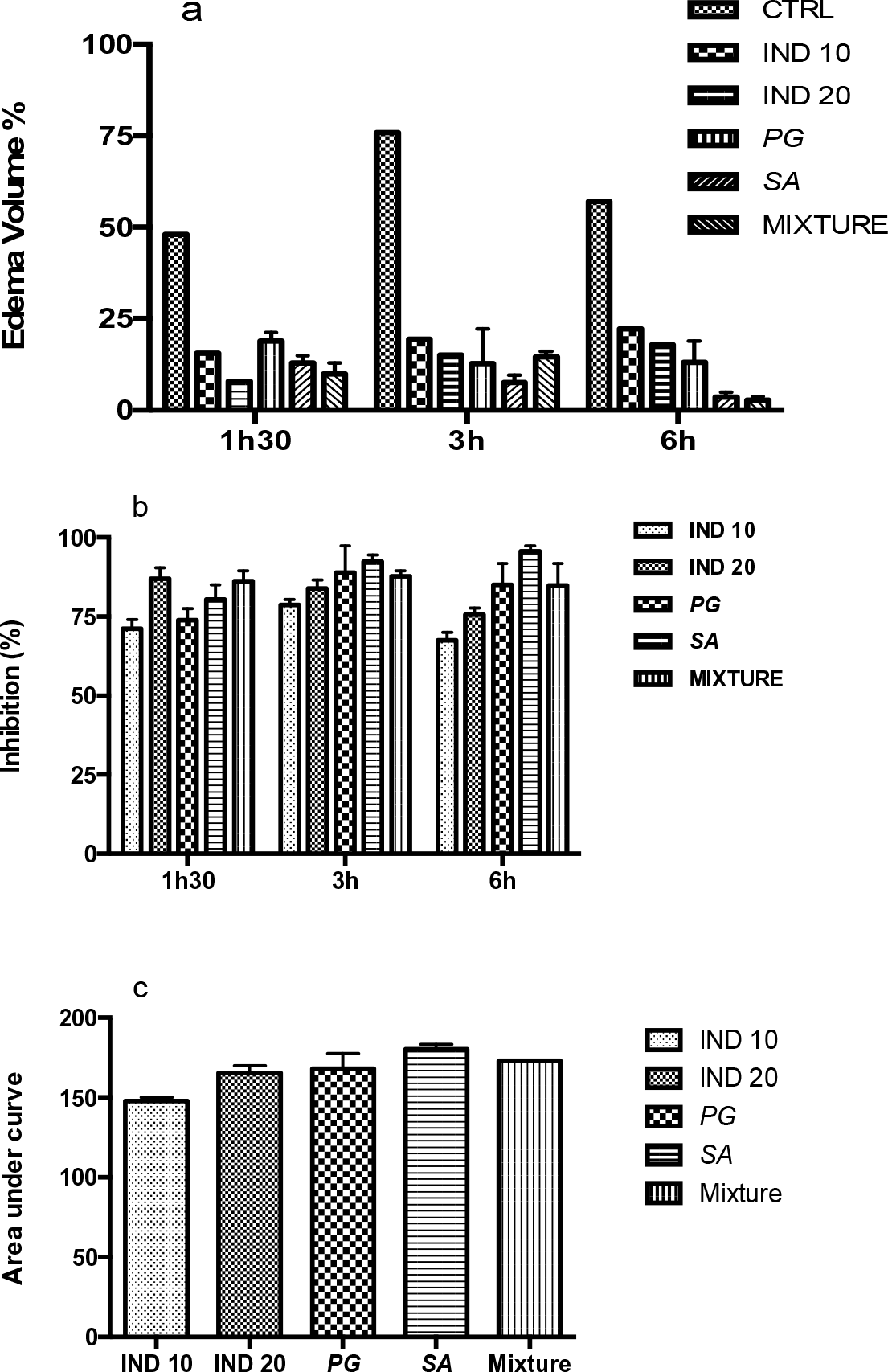
Anti-inflammatory activities. a) Edema volume. b) Inhibitory activities of EOs. c) AUC of the different treatments. Indomethacin (IND): 10 and 20 mg/Kg; EOs: 250 mg/Kg.

The variation between the PG, SA and the mixture of the both EOs can be explained mainly by their different chemical composition. Considering that the inflammatory response by carrageenan acute inflammatory model is a multi-mediated phenomenon divided into two phases: first phase (lasts up to 2 h) involves the participation of histamine, serotonin and bradykinin; the second phase (3–4 h) mainly sustained by prostaglandins and nitric oxide release [33-34]. The anti-inflammatory effects of PG, SA and the mixture of both EOs, show high potency from the 1h30 to 6th hours. Thus, EOs may therefore possess inhibitory potential at different phases of inflammation, thus making it a potent anti-inflammatory agent as shown in our results. The higher anti-inflammatory potential of PG, SA and the mixture of the both EOs, might be related to the inhibition of the release or synthesis of some mediators compounds, which might be attributed to its rich content of terpenic compounds [35], such as Eugenol, Caryophyllene, Citronellol, Geranial and 10-epi-γ-Eudesmol. The anti-inflammatory effect of these terpenic products has been previously reported [36-36-37-38], and has been demonstrated that EOs with high content of Eugenol, Citronellol, Geranial or Caryophyllene has a strong anti-inflammatory capacity against several experimental models of paw oedema [39-42]. Comparing the results on anti-inflammatory effect of SA and PG in the present work with data from previous studies, it was found that PG from Iran possesses an excellent anti-inflammatory activity [43]. Also, it has been reported previously that PG exhibited a strong anti-inflammatory effect and were effective at inhibiting proinflammatory enzymes [44]. Many authors have reported anti-inflammatory effect of SA in experimental animal models [45-46]. However, to our knowledge, the anti-inflammatory properties of EOs combinations have not been previously targeted.

## 4. Conclusion

EOs combination currently constitutes one of the most promising approaches to develop new anti-inflammatory formulations since the use of a formulation consisting of a single drug can presents serious limitations. Thus, to identify new combination compounds with anti-inflammatory properties that are more effective and less toxic compared to individual drugs seems very interesting. Several studies have shown that many drug combinations raised the pharmacological effects. For instance, a combination of several plants extracts exhibited superior activity against multiple diseases when compared to the use of individual drugs. Therefore, synergistic effect and decreased development of toxic effect of EOs are the main advantages of combination therapy. The present work demonstrated that EOs mixture of PG and SA show equivalent antioxidant and anti-inflammatory potential with lower toxicity.

## Acknowledgments

The author Ilias Marmouzi wants to thank Dr Inssaf Berkiks and Dr Taoufik Benali for their contributions. Also we are extremely thankful to Mr Abdenbi Saadi (INRA) for his valuable contribution and help.

